# Diversity of Pico- to Mesoplankton Along the 2000 km Salinity Gradient of the Baltic Sea

**DOI:** 10.1101/035485

**Authors:** Yue O.O. Hu, Bengt Karlson, Sophie Charvet, Anders F. Andersson

## Abstract

Microscopic plankton form the productive base of both marine and freshwater ecosystems and are key drivers of global biogeochemical cycles of carbon and nutrients. Plankton diversity is immense with representations from all major phyla within the three domains of life. So far, plankton monitoring has mainly been based on microscopic identification, which has limited sensitivity and reproducibility, not least because of the numerical majority of plankton being unidentifiable under the light microscope. High-throughput sequencing of taxonomic marker genes offers a means to identify taxa inaccessible by traditional methods; thus, recent studies have unveiled an extensive previously unknown diversity of plankton. Here, we conducted ultra-deep Illumina sequencing (average 10^5^ sequences/sample) of rRNA gene amplicons of surface water eukaryotic and bacterial plankton communities along a 2000 km transect following the salinity gradient of the Baltic Sea. Community composition was strongly correlated with salinity for both bacterial and eukaryotic plankton assemblages, highlighting the importance of salinity for structuring the biodiversity within this ecosystem. The distribution of major planktonic taxa followed expected patterns as observed in monitoring programs, but also novel groups to the Baltic were identified, such as relatives to the coccolithophore *Emiliana huxleyi* in the northern Baltic Sea. The deep sequencing also enabled accurate enumeration of highly resolved (> 99% identity) operational taxonomic units, which revealed contrasting distribution profiles among closely related populations, reflecting niche partitioning into ecotypes. This study provides the first ultra-deep sequencing-based survey on eukaryotic and bacterial plankton biogeography in the Baltic Sea.

## Introduction

The Baltic Sea is the world’s second largest brackish water body. It is connected to the North Sea by several narrow straits. A horizontal salinity gradient ranging from 2-30 Practical Salinity Units (PSU) traverses the Baltic Sea and parts of the North Sea in a Northeast - Southwest (NE-SW) direction. This gradient is maintained by sporadic inflows of saline deep water in the south, in combination with freshwater input from rivers. A strong halocline at 60 - 80 m depth restricts vertical mixing in the Baltic Proper, and large areas of the deeper waters are hypoxic or anoxic (Carstensen et al., 2014). The intermediate salinity and subarctic conditions constrain the macrobial biodiversity in the Baltic Sea, and its multicellular species are fewer and typically genetically impoverished compared to the populations of the NE Atlantic (Johannesson and André, 2006). In contrast, bacterioplankton diversity does not appear to decrease at intermediate salinities (Herlemann et al., 2011), and for protists it has even been suggested that species richness is actually highest in the horohalinicum region (salinity 5-8 PSU; (Telesh et al., 2011a, 2013)).

Unicellular plankton play key roles in global biogeochemical cycles and form the basis of marine food webs (Falkowski et al., 2008). In the Baltic Sea, diatoms and dinoflagellates (>10 μm) dominate primary production in spring, while smaller eukaryotic phytoplankton and cyanobacteria (<3 μm) dominate primary production over the rest of the year (Johansson, 2004). Heterotrophic bacteria play key roles in taking up dissolved organic matter stemming from primary production within the ecosystem and from terrestrial runoff-water (Azam and Malfatti, 2007). Bacteria also mediate most transformations in the cycling of nitrogen, phosphorus, trace metals and other nutrients (Falkowski et al., 2008). Energy and nutrients are propagated to higher trophic levels through grazing. Large phytoplankton are primarily grazed upon by mesozooplankton (e.g. copepods) but also by micro-zooplankton, i.e. heterotrophic protists. The current view is that picoplankton (bacteria and picoeukaryotes; 0.2-2 μm) are predated upon by nanoplankton (nanoflagellates and small ciliates; 2-20 μm). The grazing on heterotrophic bacteria brings the organic matter leaked by phytoplankton during photosynthesis back to the traditional food chain, closing what is called the microbial loop. The nanoplankton are eaten by larger zooplankton (copepods, rotifers, larger ciliates and dinoflagellates) (Samuelsson et al., 2006). Finally, the latter are predated upon by fish; in the pelagic Baltic Sea, mainly by sprat and herring. In this way, all elements of the pelagic ecosystem are interconnected, leading to cascade effects between different levels of the food web. For instance loss of top predators could be propagated all the way down to affect picoplankton community composition and function (Casini et al., 2009).

Monitoring of phyto- and zooplankton serves to estimate the environmental state and to detect changes regarding eutrophication, biodiversity, harmful algal blooms etc. This is today largely based on microscopic techniques (Hällfors, 2004; Olenina et al., 2006), which are time-consuming (~ 4 h/sample) and rely heavily on the skills of taxonomists. Cross-comparison between datasets analysed by different taxonomists may therefore be problematic. The smallest plankton organisms, i.e. the pico- and nanoplankton (0.2-2 μm and 2-20 μm), and larger organisms that lack distinctive morphological features are difficult to identify using light microscopy. Therefore, these most abundant organisms are most often simply termed ‘Unicells’ or ‘Flagellates’ since they cannot be taxonomically identified. Hence, there is a need for faster, more robust novel methods that can cover a larger fraction of planktonic diversity.

Due to the recent developments of high-throughput sequencing techniques, genetic methods are a promising avenue for monitoring biological diversity. Genetic barcoding consists of taxonomically assigning a specimen based on sequencing a short standardised DNA fragment (barcode) and by matching this to a reference library of sequences of known taxonomy. By metabarcoding, the approach is extended to a community of individuals of different species. Metabarcoding has been extensively used for assessing both prokaryotic and eukaryotic plankton diversity and biogeography (Amaral-Zettler et al., 2009; Sogin et al., 2006; de Vargas et al., 2015). A number of metabarcoding studies have investigated diversity and spatiotemporal distribution patterns of prokaryotes in the Baltic Sea (Andersson et al., 2010; Dupont et al., 2014; Herlemann et al., 2011; Lindh et al., 2015), however, the methodology has not yet been used to map eukaryotic plankton in this sea.

Here, we have conducted ultra-deep Illumina sequencing of surface-water bacterial and eukaryotic plankton communities along a 2000 km transect following the salinity gradient of the Baltic Sea. Our study, which is the first high-throughput sequencing-based investigation of eukaryotic plankton biogeography in the Baltic Sea, gives a comprehensive view of plankton diversity across the salinity gradient. It verifies previous microscopy-based knowledge on distribution patterns for many plankton groups, but also reveals taxa not previously known to exist in the Baltic.

## Materials and Methods

### Sampling

Twenty-one water samples were collected in the Kattegat, the Baltic Proper and the Gulf of Bothnia using a FerryBox system installed in the ship TransPaper during 13^th^-19^th^ of July 2013. The ship followed the route: Gothenburg (Sweden) - Kemi (Finland) - Oulu (Finland) -Lübeck (Germany) - Gothenburg. The FerryBox system consists of a pump with a water inlet at 3 m depth, a circuit of multiple sensors for temperature, conductivity, chlorophyll and phycocyanin fluorescence, turbidity and oxygen as well as automated water sampling devices. A detailed description of the FerryBox system is found in Karlson et al. (submitted). Manual water sampling for DNA analysis was carried out both on the Northward and Southward legs. Approximately 10 L of seawater were collected in a polycarbonate carboy. Subsamples of 200 to 500 mL were filtered onto 0.22 μm pore-size mixed cellulose ester membrane filters (Merck Millipore co., Cat. No. GSWP04700) to capture plankton. The filters were frozen in liquid nitrogen on board and kept at -20 °C to -80 °C until DNA extraction.

### DNA Extraction, PCR Amplification and Sequencing

Genomic DNA was extracted using the PowerWater^®^ DNA isolation kit (MO-BIO Laboratories Inc, Carlsbad CA, USA) following the instructions provided by the manufacturer. The V3-V4 regions of bacterial 16S rDNA were PCR amplified with primers 341F (CCTACGGGNGGCWGCAG) and 805R (GACTACHVGGGTATCTAATCC) (Herlemann et al., 2011), and the V4-V5 regions of eukaryotic 18S rDNA were amplified with primers 574*F (CGGTAAYTCCAGCTCYV and 1132R (CCGTCAATTHCTTYAAR) (Hugerth et al., 2014a)), using KAPA HiFi HotStart ReadyMix (2X) (KAPA Biosystems, Kit Code KK2602). A two step PCR procedure was applied (Hugerth et al., 2014a), with 35 and 38 PCR cycles in total for 16S and 18S rDNA, respectively. Between the first and second PCR, and prior to pooling libraries, amplicons were purified with 8.8% PEG 6000 (Polyethylene Glycol 6000) (Merck Millipore co., Cat. No. 528877-1KG) precipitation buffer and CA beads (carboxylic acid-coated superparamagnetic beads) (Dynabeads^®^ MyOne™ Carboxylic Acid, Cat. No. 65012)(Lundin et al., 2010). Agilent 2100 Bioanalyzer (Agilent, Technologies, DNA 1000 LabChip kit) and Qubit^®^ 2.0 Fluorometer (Invitrogen, Qubit-IT™ dsDNA HS Assay kit) were used for checking the amplicon fragment sizes and quantification. Equimolar amounts of indexed samples were mixed and sequenced with Illumina MiSeq (Illumina Inc, USA) at NGI/Scilifelab Stockholm. The sequencing reads have been submitted to the European Nucleotide Archive (ENA) under accession numbers XXXXXX.

### Sequence Processing

The OTU table building procedures followed https://github.com/EnvGen/Tutorials, using USEARCH for quality trimming and OTU-clustering (Edgar, 2010), but with some optimizations. The primers were trimmed from forward and reverse reads with FastX (Pearson et al., 1997), as well as 40 bases in the 3’ end of the reverse reads (these displayed lower quality than the forward reads). This resulted in average Phred scores (Q values) of every position in the trimmed reads to be >20. In order to avoid generating artificial OTUs while assigning as many reads as possible, we used more stringent criteria when generating OTU-centroid sequences than in the mapping (assigning step). For the centroid generation step, 3’-ends were further trimmed such that all remaining bases had Q values ≥ 25. Forward and reverse reads were merged, and merged pairs that were shorter than 300 bp, had > 3 mismatches, or overlapped by < 100 bp were discarded. Remaining merged reads were clustered into OTUs using 99% identity level. For processing reads for the assigning step, the same procedure was applied, but with Q20 as the quality trimming parameter. The trimmed reads were mapped to the OTU centroids to build the 16S OTU table. 16S rDNA OTUs (their centroid sequences) were classified with SINA-1.2.13 using the SILVA 119 SSU database (Quast et al., 2013).

The 18S OTU table was built similarly to the 16S OTU table, but the overlap between read pairs was not sufficient for merging, so each pair was concatenated rather than merged. Primers were trimmed from forward and reverse reads, as well as 44 bases in the 3’ end of the reverse reads, resulting in average Phred scores of every position to be > 20. After trimming, forward and reverse reads were concatenated prior to OTU clustering. For the taxonomic classification, OTU centroid sequences were split at the concatenation position, and the forward and reverse counterparts were separately BLAST (Altschul et al., 1997) searched against the PR2 database (based on GenBank 203 - October 2014)(Guillou et al., 2013). BLAST results were filtered such that only alignments of at least 250 and 210 bp in length (90% of the length of reads), for the forward and reverse read, respectively, were kept. Then, the highest taxonomic level was found where the forward and reverse read aligned to the same database sequence, with alignment identities exceeding the taxonomic level-specific cutoff. If just one database sequence was hit, the OTU inherited the taxonomy of this sequence. If multiple database sequences were hit, the bit-scores of the forward and reverse reads to each hit were summed, and the hits having at least 95% of the best sum of bit scores were selected, and the most detailed consensus taxonomy was built from these hits and assigned to the OTU. The taxonomic-level specific identity cutoffs were as follows: kingdom to family level (PR2 level 1 - 6): 90%; genus level (PR2 level 7): 97%, species level (PR2 level 8): 99%. These cutoffs were determined by running an in-silico experiment where artificial reads were generated from 1000 randomly selected PR2 sequences, and classified against the remaining sequences in PR2, using a range of identity cutoffs at each taxonomic level. The selected cutoffs were good compromises between percentage of sequences classified and classification accuracy. In addition to this automatic taxonomic annotation procedure, the taxonomy was manually determined for a subset of the individual OTUs mentioned in the text by online BLAST searches against the NCBI nt database.

### Statistical Analysis

All statistical analysis and plotting were conducted in R (www.r-project.org) using the R libraries: vegan (α-diversity; β-diversity), rworldmap (geographic plotting), ape (pcoa), cluster (hierarchical clustering), pheatmap (heatmap).

## Results

### Trends in Beta-diversity

On average 32,948 amplicons of bacterial (16S) and eukaryotic (18S) rRNA genes were sequenced from surface water samples from 21 stations within the Baltic Sea and Kattegat, collected in July 2013 (Figure 1A). The sequences were clustered into 1269 and 2022 16S and 18S operational taxonomic units (OTUs), respectively. To investigate overall trends in community composition, Principal Coordinates Analysis (PCoA) was conducted on the bacterial and eukaryotic communities (Figure 1). Similar to what has been shown before in the Baltic Sea (Dupont et al., 2014; Herlemann et al., 2011), bacterial community composition changed gradually along the salinity gradient, with the first principal coordinate being highly correlated with salinity (Spearman rho = 0.99, P < 10^−5^; Figure 1B). Here we could also, for the first time, show a similar pattern for eukaryotic plankton in the Baltic (Spearman rho = 0.88, P < 10^−5^; Figure 1C). Likewise, hierarchical clustering grouped the samples according to their salinity, based on both bacterial and eukaryotic community composition (Supplementary figure 1). Difference in salinity between samples was also directly correlated with difference in community composition (beta-diversity), for both bacterial and eukaryotic communities (Supplementary figure 2). Difference in temperature, that to some extent covaries with salinity (Supplementary table 1), was also correlated with beta-diversity, but the correlation was weaker (Supplementary figure 2). Since both eukaryotic and bacterial communities correlated with salinity, it followed that beta-diversity of the two community types were correlated (Figure 1D).

**Figure 1.**
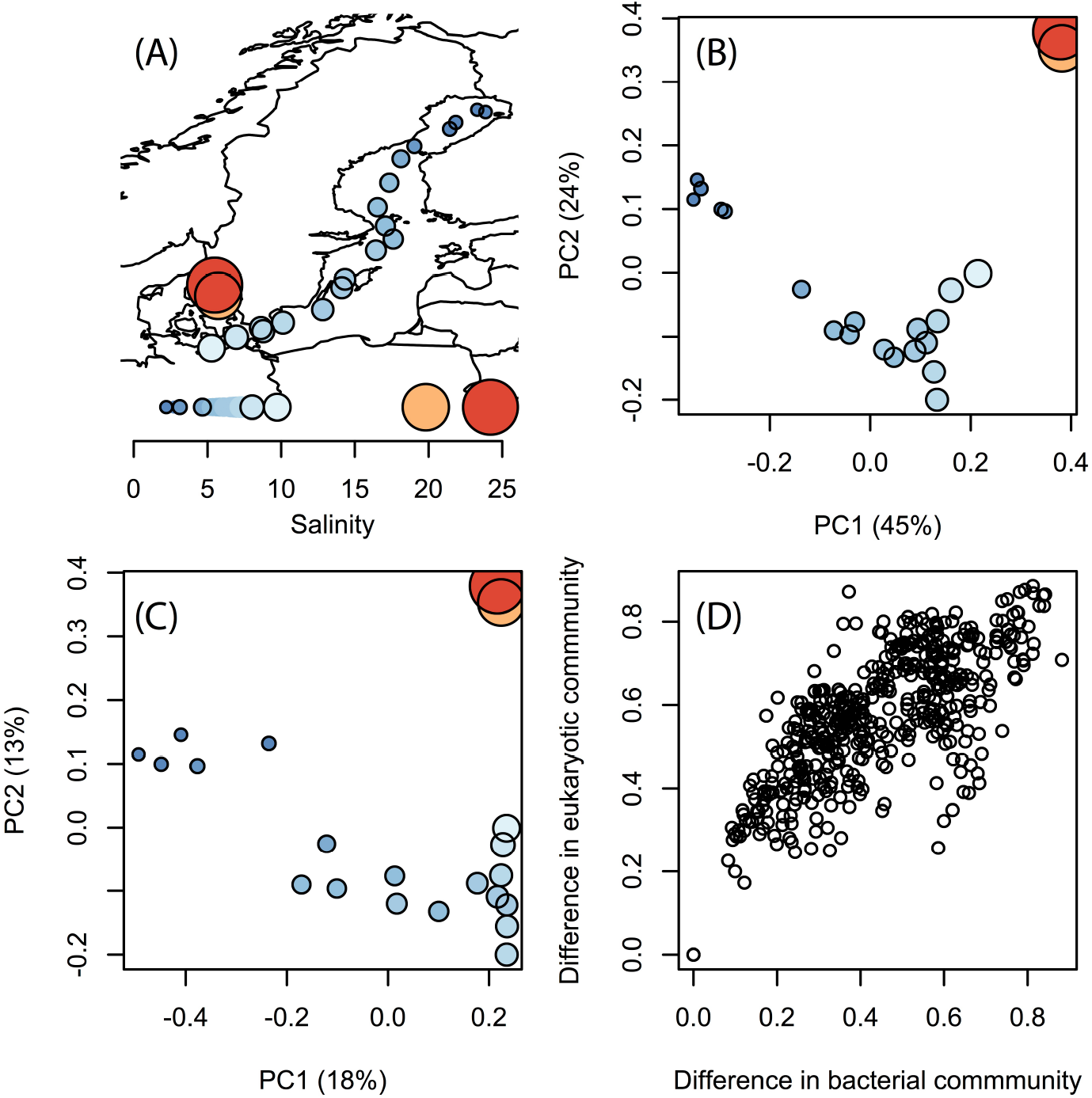
Trends in bacterial and eukaryotic beta-diversity. (**A**) Map showing sampling sites within Kattegat and the Baltic Sea, with salinity indicated by size and color. **(B-C)** Ordination of bacterial (B) and eukaryotic (C) communities using Principal Coordinate Analysis (PCoA) based on beta-diversity calculated using Spearman correlation of OTU frequencies. Samples are colored and sized according to (A). Variation explained by the principal components (PCs) are given within parenthesis. PC1 was highly correlated with salinity for both bacterial and eukaryotic communities (Spearman *ρ* > 0.97 and P < 10^−5^ for both). **(D)** Correlation of beta-diversity in bacterial and eukaryotic communities (each circle is one pair of samples).

**Figure 2.**
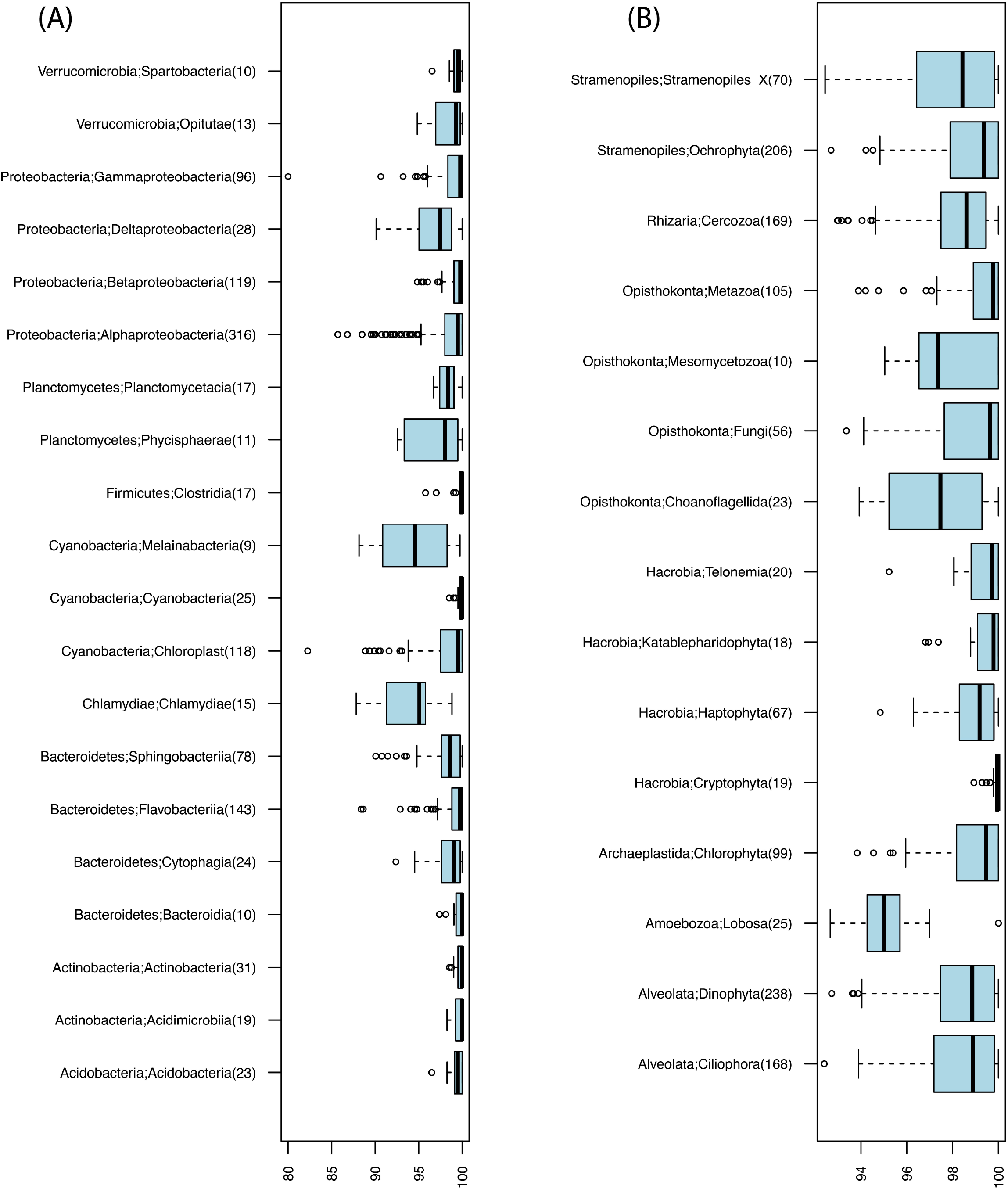
Identity levels (%) between OTU centroid sequences and their best matching database sequence. (**A**) Bacterial OTUs. **(B)** Eukaryotic OTUs. Number of OTUs included in each taxonomic group is given within parenthesis; only groups with ≥10 OTUs are shown.

### Bacterial Community Composition

Our analysis confirmed the overall trends in bacterial taxonomic composition observed in previous studies, with an increase in Alpha- and Gammaproteobacteria and a decrease in Actinobacteria and Betaproteobacteria with increasing salinity levels (Dupont et al., 2014; Herlemann et al., 2011). Similar to what was found by Herlemann et al. (Herlemann et al., 2011), Verrucomicrobia peaked in intermediate salinity levels, which here was also the case for Planctomycetes and Cyanobacteria (Figure 3A).

**Figure 3.**
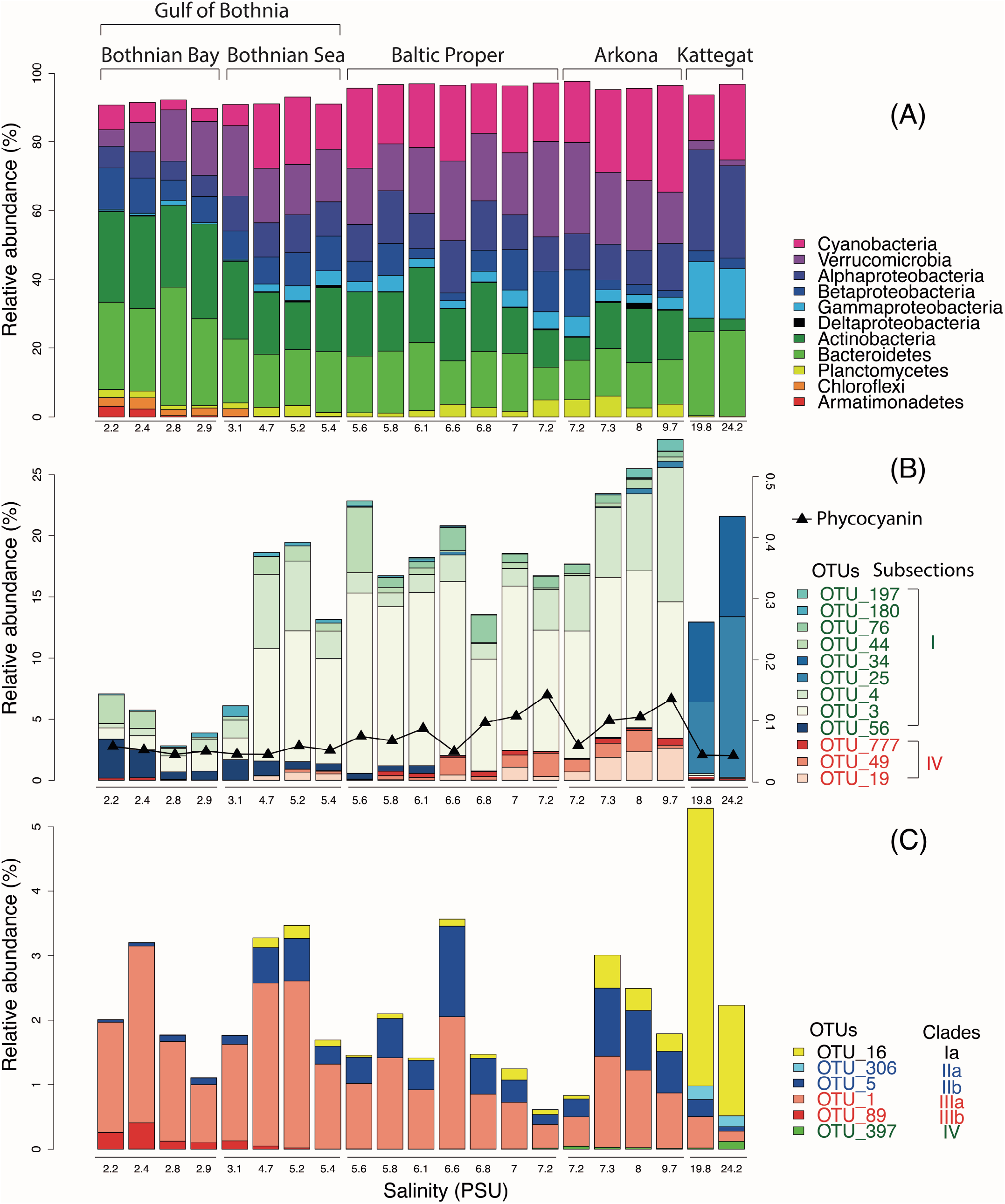
Abundance distribution of major bacterial groups across the Baltic Sea salinity gradient. **(A)** Major bacterial taxonomic groups (>0.1% mean abundance). **(B)** Dominant Cyanobacteria OTUs (> 0.1% mean abundance). In Subsection I, OTU_56 is a *Microcystis* (a freshwater species), remaining OTUs belong to *Synechococcus*. In Subsection IV, OTU_777, OTU_49 and OTU_19 belong to *Aphanizomenon*, *Nodularia* and *Dolichospermum* (syn. *Anabaena*), respectively. The black curve indicates phycocyanin fluorescence (in arbitrary units). **(C)** Dominant SAR11 OTUs (> 0.1% mean abundance).

Of the 1243 16S OTUs observed in the 21 samples, 1187 could be taxonomically classified at least to the class level. The majority (59%) of the OTU centroid sequences matched SILVA database reference sequences with > 99% identity. Identity levels however varied substantially among taxa (Figure 2). Chlamydiae and the newly discovered non-photosynthetic sister clade to Cyanobacteria, the Melainibacteria (Di Rienzi et al., 2013), displayed notably low identity levels (Figure 2A). Aquatic Chlamydiae are symbionts on e.g. fish and amoeba (Collingro et al., 2005; Stride et al., 2014) and likely difficult to be isolated and cultured, and likewise no isolates have yet been obtained for the Melainibacteria. The Deltaproteobacteria, that are abundant in deeper waters of the Baltic Sea (Herlemann et al., 2011), also displayed comparatively low identity levels.

The gradual shift in community composition observed at the phylum/class level was also evident within specific taxonomic groups, as exemplified by the Cyanobacteria and the Alphaproteobacteria cluster SAR11 in Figure 3 B,C. The Cyanobacteria were represented by 26 OTUs, of which twelve had a mean abundance of >0.1% of reads, most of which were related to the picocyanobacteria *Synechococcus* (Figure 3B). OTUs associated to filamentous cyanobacteria (*Aphanizomenon*, *Nodularia* and *Dolichospermum* (syn. *Anabaena*)) were also detected from the brackish region but in much lower abundances than the dominant *Synechococcus*. Phycocyanin fluorescence, as recorded by the FerryBox system, correlated well with relative abundance of sequenced filamentous cyanobacteria (Spearman rho = 0.69, P < 10^−3^), in agreement with Seppälä et al that observed a linear relation between biomass of filamentous cyanobacteria and phycocyanin fluorescence (Seppälä et al., 2007). While massive blooms of filamentous cyanobacteria in the summer are characteristic of the Baltic Sea, the majority of cyanobacterial biomass and cell numbers are in fact usually comprised of picocyanobacteria (Stal et al., 2003). The different *Synechococcus* populations showed clear differences in salinity preference, with OTU_25 and OTU_34 being dominant in Kattegat at more than 99.7% of cyanobacterial reads, while OTU_3 and OTU_4 dominated the Baltic Proper with on average 88% of cyanobacterial reads.

SAR11 is the most abundant bacterioplankton clade of the world’s oceans, and is also found in brackish and fresh waters. It consists of several subclades that together span the candidate order Pelagibacterales (Grote et al., 2012). Eight OTUs were classified as SAR11 in our dataset, of which six had >10 reads (Figure 3C), representing five different subclades of SAR11 (Supplementary figure 3 for a phylogenetic tree). An OTU identical to *Pelagibacter ubique* (open ocean subclade Ia) dominated the marine samples but was also present in the brackish samples, while two OTUs belonging to subclade II and IIIa dominated the brackish region. Subclade IIIa is frequent in brackish waters and has previously been found in the Baltic Sea (Bergen et al., 2014; Dupont et al., 2014; Herlemann et al., 2011), while subclade II is mainly associated with coastal and mesopelagic waters (Carlson et al., 2009; Michael Beman et al., 2011). Interestingly, contrary to previous studies (Bergen et al., 2014; Dupont et al., 2014), we also found an OTU representing the freshwater subclade IIIb (LD12) in the low-salinity Bothnian Bay.

The deep sequencing provided by the Illumina platform allowed clustering of reads with higher phylogenetic resolution than with earlier 454-data (99% identity; compared to 97% in the study by Herlemann (Herlemann et al., 2011)) while still maintaining enough reads to accurately quantify OTUs. This uncovered highly similar OTUs with distinct spatial distributions. In the study by Herlemann et al. (2011), the most abundant OTU was a Verrucomicrobia belonging to the class Spartobacteria, for which the genome was later reconstructed with metagenomics (Herlemann et al., 2013), and for which the name Candidatus *Spartobacteria baltica* was proposed. In the current dataset we obtained an OTU identical to *S. baltica* (OTU_2) but we also obtained closely related OTUs (>98% identity to OTU_2) with contrasting distribution patterns over the salinity gradient (Supplementary figure 4).

### Eukaryotic Community Composition

We retrieved 18S rRNA gene reads corresponding to nine eukaryotic superphyla, 24 phyla, 77 classes and 256 genera (see Figure 5 for an overview). Since no prefiltration was applied before the plankton were captured on filters, the eukaryotic organisms span a large size range, and include unicellular organisms (protists) as well as multicellular organisms (metazoa). The phyla Metazoa, Ciliophora, Dinophyta, Cercozoa, Chlorophyta, Katablepharidophyta, Stramenopiles_X, Ochrophyta, Telonemia, Haptophyta and Cryptophyta each makes up >1%, and together comprise 94% of the total 18S sequence reads. Stramenopiles_X is a division used in PR2 and includes heterotrophic unicellular heterokonts, e.g. the Oomycota, MAST and Labyrinthula. Some samples were highly dominated by metazoan sequences (mainly copepods, Figure 4C), likely due to the large number of rRNA genes in their genomes and the fact that these are multicellular organisms. Nevertheless, they never exceeded 70% of the sequences in a sample, leaving at least 17,226 sequences for description of the remaining community of unicellular protists.

**Figure 4.**
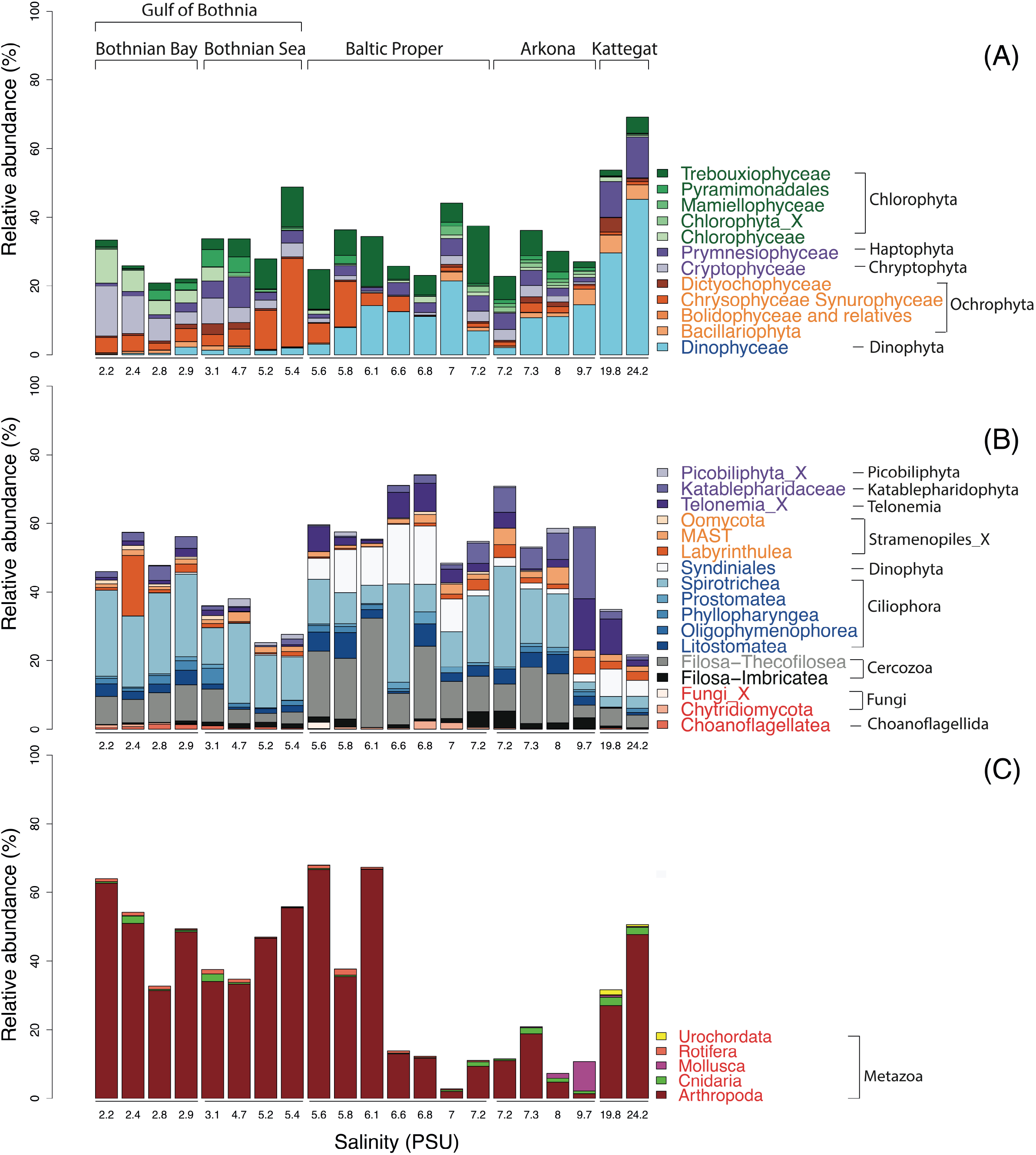
Abundance distribution of major eukaryotic taxonomic classes of plankton across the Baltic Sea salinity gradient. Classes displaying >1% mean abundance are shown and colored according to the superphylum they belong to. Green, purple, orange, blue, grey and red represent Archaeplastida, Hacrobia, Stramenopiles (or Heterokonta), Alveolata, Rhizaria and Opisthokonta, respectively. **(A)** Phytoplankton, here broadly defined as groups of organisms that include mainly phototrophic and mixotrophic representatives, although some groups, such as Dinophyta, also include organisms that are heterotrophic. **(B)** Heterotrophic plankton excluding metazoa. **(C)** Metazoa. Relative abundnaces were in (A) and (B) calculated as percentages of protist (i.e. non-metazoan) sequence reads; in (C) as percentages of all eukaryotic sequence reads.

**Figure 5.**
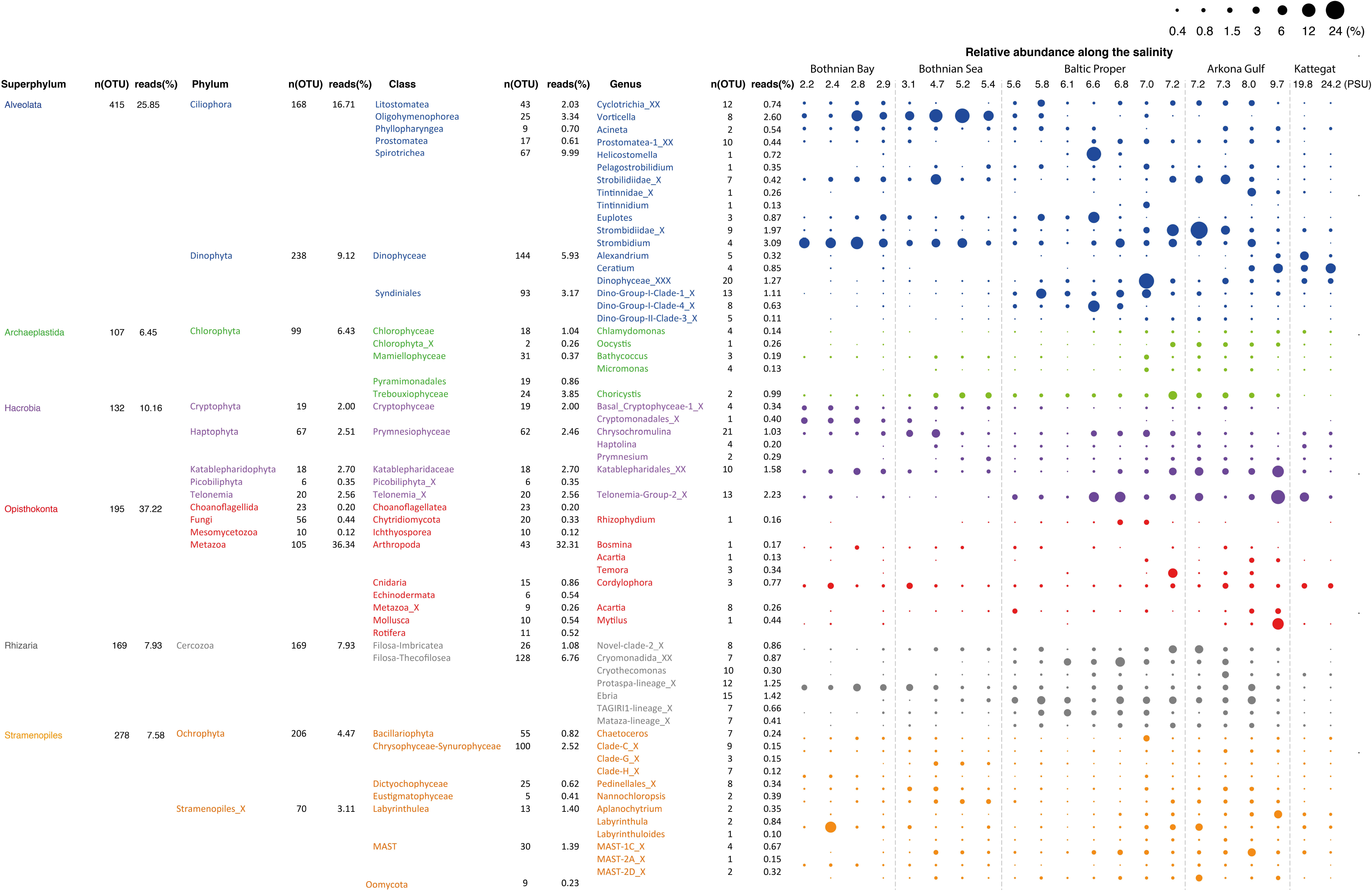
Summary of detected eukaryotic taxonomic groups. The number of detected OTUs and the mean abundance is given for each taxonomic group at the superphylum, phylum, class and genus level (corresponding to PR2 levels 2, 3, 4 and 7). Only groups displaying >0.1% mean abundance are shown. The dot plot illustrates the distribution of the annotated genera along the salinity gradient, with circle area corresponding to relative abundance.

Of the total 1860 18S OTUs, 283 could be attributed to a known genus or species, 309 could be assigned to PR2 phylogenetic clusters corresponding to the genera level, and 740 could only be classified to higher taxonomic levels. Remaining OTUs could not be classified using the PR2 database, but these corresponded to only 5.63% of the reads in the samples on average. As was observed for the bacteria, identities of closest-related database sequences varied substantially between clades (Fig. 2B). Groups with unusually low similarity levels (median < 98%) were Amoebozoa, Choanoflagellida, and Mesomycetozoa. The particularly low identity levels of Amoebozoa and their potential bacterial symbionts, Chlamydiae (see above), thus indicate that the Baltic Sea is inhabited by a set of poorly characterised Amoebozoa harbouring poorly characterised bacterial symbionts. The Mesomycetozoa (syn. Ichthyosporea) are deficiently described fungus-like protists known to live in association with a wide range of host organisms (Glockling et al., 2013).

### Eukaryotic Phytoplankton

Recognizing that some protistan groups include phototrophic, mixotrophic and heterotrophic organisms, a division of the organisms in phytoplankton and heterotrophic plankton has been made. Here mixotrophs are treated as the phototrophic phytoplankton. Among the dinoflagellates many are mixotrophic, i.e. they carry out photosynthesis but also feed on other organisms.

Microphytoplankton, made up of diatoms and dinoflagellates, are characteristic of the saline southern and western parts of the Baltic Sea and Kattegat, while smaller eukaryotic phytoplankton groups are more dominant in the northern Baltic Sea (Ojaveer et al., 2010). Pronounced seasonal fluctuations also prevail, with diatoms and dinoflagellates blooming in spring, and smaller eukaryotic phytoplankton and cyanobacteria dominating in summer (Johansson et al., 2004). Diatoms (Bacillariophyceae) and dinoflagellates (Dinophyceae) had higher relative abundances in the more saline regions of our survey (Figure 4A). However, although the samples were taken after the spring bloom, a substantial fraction of the reads from the Baltic Proper also broadly classified as dinoflagellates (Figure 5A). On average, dinoflagellates contributed to 10% of the reads. The relative proportion of dinoflagellates decreased from 50% to nearly 0% of the protist fraction, going from highest to lowest salinity (Figure 4A).

In the high salinity Kattegat, dominant dinoflagellate genera were *Alexandrium* and *Ceratium* of the order Gonyaulacales (Supplementary figure 5B), while in the Baltic Proper, typical brackish dinoflagellates from the order Suessiales, such as *Biecheleria baltica* (OTU_25), were most abundant. Major OTUs of diatoms belonged to species often found in Baltic monitoring programs, such as the polar centric diatoms *Attheya longicornis* (OTU_144) and *Chaetoceros neogracile* (OTU_136), and the raphid diatom *Cylindrotheca closterium* (OTU_94). Interestingly, at the station south of Öland in the Baltic Proper (salinity 6.97 PSU), the Bacillariophyta and Dinophyceae were both dominated by OTUs (OTU_125 and OTU_32) that displayed highest similarity (99.7% and 98.1% identity) to sequences of uncultured diatoms and dinoflagellates from anoxic layers of the Framvaran Fjord in Norway ((Behnke et al., 2010); Supplementary figure 5). Why these potentially anaerobic diatoms and dinoflagellates flourished in the surface water at this particular station remains to be explained.

Unlike the distribution pattern of dinoflagellates and diatoms, other phytoplankton, such as Chrysophyta, Chlorophyta and Cryptophyta, were observed at lower salinities. The typical fresh-water groups, Chrysophyceae-Synurophyceae (these two classes are combined in the PR2 database), were abundant in the surface water in the Gulf of Bothnia, peaking with 25% of protist reads at 5.4 PSU in the Bothnian Sea (Figure 4A). The dominant OTU (OTU_4) in this area belonged to *Uroglena* (100% identity), a common colony-forming genus, that made up nearly half of the phototrophic protist community reads. The green algae group, Chlorophyta, was found prevalent in the Baltic Proper and Gulf of Bothnia (Figure 4A, Figure 5). In the Baltic Proper it was dominated by the genus *Choricystis* (class Trebouxiophyceae) (OTU_9), while in the Gulf of Bothnia two Chlorophyta freshwater OTUs (OTU_103: order Chlamydomonadales, OTU_65: *Monoraphidium convolutum*), together with one Cryptophyta freshwater species (OTU_56: *Rhodomonas minuta*) made up one third of the phytoplankton reads (Supplementary figure 5C).

The ultra-deep sequencing also revealed organisms not easily identifiable with a microscope, such as picoeukaryotes of the class Mamiellophyceae within the Chlorophyta (Figure 5), from which we detected OTUs of the genera *Ostreococcus, Micromonas* and *Bathycoccus* (Supplementary figure 5E). Though they never contributed with more than 2.5% of the protist reads, they displayed distinct distribution patterns along the salinity gradient. For example, *Ostreococcus*, the smallest known eukaryote, was associated with two OTUs that were separately abundant in Bothnian Sea (OTU_315) and Arkona Basin (OTU_464). Notably, the OTU that peaked in the Bothnian Sea was absent from the high salinity Kattegat region. This OTU belonged to *Ostreococcus* clade D according to its high similarity (1 deletion) to clade D strains isolated from the Mediterranean (Subirana et al., 2013), while OUT_464 belonged to clade C with its representative sequence being identical to 18S of the widespread *Ostreococcus tauri* (Keeling, 2007). Our results were consistent with a study on global distribution of *Ostreococcus* (Demir-Hilton et al., 2011), which reported that Clade D was only found in estuarine or brackish waters, while Clade C was also found in marine waters.

The most abundant Haptophyta OTUs were most closely related to Coccolithophorids (class Prymnesiophyceae), which are globally important phytoplankton with characteristic exoskeletons of calcium carbonate plates (coccoliths). They are found in large numbers throughout the ocean’s euphotic zone but only one species, *Balaniger balticus*, has been described in the Baltic Sea (Thomsen and Oates, 1978). The sparsity of coccolithophorids in the Baltic has been attributed to undersaturation of calcium carbonate in winter, potentially resulting in dissolution of the coccoliths (Tyrrell et al., 2008). We found two coccolithophore-related OTUs (OTU_127 and OUT_2007) that reached a maximum of 4.2% of the protist reads in the high salinity samples in Kattegat. The representative sequences of these two OTUs showed 100% identity to both coccolithophore species *Emiliania huxleyi* and *Gephyrocapsa muellerae* from the family Noelaerhabdaceae. Interestingly, these two species have identical sequences (Bendif et al., 2015). Intriguingly, OUT_127 was also detected, though at much lower abundances, in the Gulf of Bothnia (Supplementary Figure 6). Here we also detected the non-calcifier *Isochrysis galbana* (OTU_585) that belong to the same order (Isochrysidales) as the coccolithophores. As far as we know, neither *Emilianial/Gephyrocapsa* or *Isochrysis* have been observed in the Baltic Sea before. Although OTU_127 dominated the prymnesiophytes in Kattegat, coccolithophores were not the predominant organisms from this group in the Baltic regions. Instead, order Prymnesiales was dominant with OTUs widely spread according to their salinity preferences (Figure 5). *Prymnesium*, a genus that include the potentially toxic spring-bloom forming species *Prymnesium polylepis (Gorokhova et al., 2014)*, was present in all samples except those collected in the Bothnian Bay (Figure 5).

While the above data on eukaryotic phytoplankton are based on 18S rRNA gene amplicons, phytoplankton can potentially also be monitored using 16S sequencing of chloroplast sequences (Eiler et al., 2013; Lemieux et al., 2014). Since our 16S primer pair match a fair amount of chloroplast sequences (65% of chloroplast sequences in RDP v.11.4 (Cole et al., 2014)), 120 of the 16S OTUs, corresponding to on average 0.3% of the reads, were classified as chloroplasts by SILVA, and a subset of these (98 OTUs) could be further classified by PR2. Ideally, the 16S and 18S data should reveal the same trends across samples for the same clade or species. We tested this by first correlating relative counts at the class level. For the seven classes that had at least 1000 reads in total in each dataset (16S and 18S), six displayed significant correlations (P < 0.05; average Spearman correlation coefficient = 0.75). Comparing at a more detailed taxonomic level was complicated by the fact that the 16S OTUs lacked a detailed taxonomic annotation. Instead we correlated all 16S OTUs to all 18S OTUs and extracted the pairs displaying a Spearman rank order correlation coefficient of at least 0.8. Of the 304 pairs found, six included a 16S OTU classified as chloroplast. For five of these six pairs, the 16S and 18S OTU were classified to the same taxonomic class (four different classes) (Supplementary figure 7). The probability that this level of matching would occur by chance is extremely low, and hence these five pairs of OTUs likely represent the same species or genus. This demonstrates the robustness of the metabarcoding approach and indicates that this type of correlation analysis can be fruitful for linking chloroplast 16S sequences to their nuclear 18S counterparts.

### Eukaryotic Heterotrophs

The detected heterotrophic protists mainly belonged to the phyla Ciliophora, Cercozoa, Dinophyta, Katablepharidophyta and Telonemia (Figure 4B). Ciliates (Ciliophora) that carry out photosynthesis by using klepto-chloroplasts belong to the functional group phytoplankton but are treated among the heterotrophs is in this article. Heterotroph dominance varied between class Spirotrichea from the phylum Ciliophora, in the Gulf of Bothnia and the Southern Baltic, and class Filosa-Thecofilosea from the phylum Cercozoa, in the Northern Baltic Proper (Figure 4B). The abundance of both these classes dropped in the Kattegat (19.8 and 24.2 PSU), as well as in the “intermediate” Arkona sample (9.7 PSU).

Of the 173 OTUs classified as Ciliophora, 32 had a mean relative abundance >10^−3^. The most prominent OTUs belonged to the genus *Vorticella* (OTU_2) from class Oligohymenophorea and to *Strombidium* (OTU_5) from class Spirotrichea (Figure 5, Supplementary figure 5F). Only two OTUs (OTU_425, OTU_666), that displayed rather low abundance, belonged to the genus *Mesodinium(Mironova et al., 2009)*, normally abundant in the Baltic Sea. The low yields could be due to diurnal vertical migration, or that the *Mesodinium* population was present in thin layers that were not sampled (Sjöqvist and Lindholm, 2011). Cercozoa is a highly cryptic phylum that contains widespread protozoan omnivores and parasites with vague morphological characters (Bass et al., 2009; Cavalier-Smith, 1998; Weber et al., 2012). Of the 170 Cercozoa OTUs, 25 had a mean relative abundance >10^−3^ (Figure 5, Supplementary figure 5G).

Other heterotrophic strategies, such as parasitism, also play essential roles in the microbial food web, but monitoring parasites or symbionts by microscopy can be difficult. Sequencing of marker genes can greatly facilitate depiction of the distribution pattern of these organisms. For instance, Syndiniales, a class of Dinophyta that are parasites on other dinoflagellates, were found where other dinoflagellates were abundant (Fig 4 A,B). In total, 93 OTUs were found from three Syndiniales groups (group I, II, III) (Guillou et al., 2008). The dominant OTUs were mainly from group I (Figure 5), while group II was the most diverse with 56 OTUs.

In the Hacrobia division, three major heterotrophic classes, Picobiliphyta, Katablepharidaceae and Telonemia, showed contrasting distribution patterns in the Baltic. Initially described from cold polar waters (Lovejoy et al., 2006), Picobiliphyta recently renamed Picozoa (Seenivasan et al., 2013) is a highly diverse deep-branching clade, with representatives found in a variety of marine ecosystems (Not et al., 2007, 2009). In the present study, the dominant Picobiliphyta OTU (OTU_44), possessing almost 80% of the Picobiliphyta reads, was found identical to the only cultured member of the Picozoa, *Picomonas judraskeda* (Seenivasan et al., 2013). These picoeukaryotes (2-5 μm length) have previously been enriched from brackish Baltic Sea water under dark incubations (Weber et al., 2012). Abundance of Picobiliphyta peaked in the Bothnian Sea, while the larger Katablepharydophyta and Telonemia accumulated mainly in the Baltic Proper and the Arkona basin. Besides salinity, their distinct feeding habits are also a potential factor underlying the observed distributions. The Katablepharidaceae and Telonemia feed on prey of overlapping sizes, principally bacteria and small phytoplanktonic cells, whereas *Picomonas judraskeda* gains energy by uptake of small organic particles (Seenivasan et al., 2013).

The superphylum Stramenopiles were represented by the Ochrophyta and Stramenopiles_X. The 70 Stramenopiles_X OTUs were mainly annotated to the classes Labyrinthulea, Oomycetes and MAST. One of the samples from Bothnian Bay (salinity 2.4 PSU) displayed markedly higher abundance of Labyrinthulea than the others (Figure 4B). Class Labyrinthulea was highly dominated by an OTU (OTU_54) that was found in all Baltic regions with no apparent salinity preference. The representative sequence of OTU_54 was highly similar (99 - 100% identity) to endophytic Labyrinthula strains isolated from either mangrove leaves or eelgrass, which indicates that OTU_54 is an endophyte associated with plants. Unlike other sampling sites, the depth at this station is rather shallow (17-meters), and directly to the east a region of shallow waters (10-meters depth on average) extends to the Finnish coast. Hence, the high abundance of Labyrinthula here could be due to debris from benthic macro vegetation captured on the water filter.

Of the 70 OTUs from Stramenopiles_X, 30 were annotated to the MAST lineage, a diverse group of pico- to micro-sized heterotrophic marine stramenopiles that was discovered by environmental sequencing only ten years ago (Massana et al., 2004) and has not yet been cultured (Massana et al., 2006). Many studies have since then investigated the links between environmental factors and MAST lineages with DNA sequencing (Massana et al., 2004, 2014) and fluorescent in situ hybridization (FISH) (Lin et al., 2012; Piwosz and Pernthaler, 2010; Thaler and Lovejoy, 2013). In this study, seven MAST clades (“ribogroups”; (Massana et al., 2014)) were detected. The most prevalent ones were MAST-1 and -2 (Figure 5, Supplementary figure 5H), which are commonly found in marine and coastal regions (Massana et al., 2004, 2006). To date, four subclades of MAST-1 are known (Massana et al., 2014), however, only MAST-1C was observed in our dataset, with five associated OTUs (Supplementary figure 5H). Besides the most prevalent MAST clades (MAST-1C, -2D), MAST-4D and -6 were abundant in the brackish region (4 - 10 PSU). For MAST-6, all OTUs (OTU_145, 245 and 644) were missing in Bothnian Bay and Kattegat, which indicates these are brackish specialists. In the Gulf of Bothnia, MAST-2A, -3J, -12C and -12_X displayed pronounced preference of low salinity (<4 PSU) (Supplementary figure 5H). So far, only sequences from MAST-2 and -12 have been found in freshwater (Massana et al., 2014). The OTU (OTU_337) from MAST-3J that only appeared in the Bothnian Bay indicates that also this subclade contains freshwater members. In contrast, other MAST-3 subclades only appeared in the high salinity Kattegat samples.

The detected unikonts (superphylum Opisthokonta) in this study included Choanoflagellates, Mesomycetozoa, Fungi and Metazoa, with 195 OTUs annotated in total. The most divergent and abundant group was Metazoa with 105 OTUs of which 15 had a mean relative abundance >10^−3^. A recent review of plankton diversity in the Baltic ranked 23 abundant mesozooplankton species in different subbasins based on monitoring data (Ojaveer et al., 2010). OTUs corresponding to these Metazoa species displayed remarkably similar distribution patterns (Figure 6). Unicellular unikonts (Choanoflagellates, Mesomycetozoa, Fungi) made up a limited fraction of protist reads (Figure 4B). The low abundance of choanoflagellates may be attributed to seasonal dynamics, since they have earlier been reported to, unlike other heterotrophic nanoflagellates, be sparse during summer in the Baltic Sea (Samuelsson et al., 2006).

**Figure 6.**
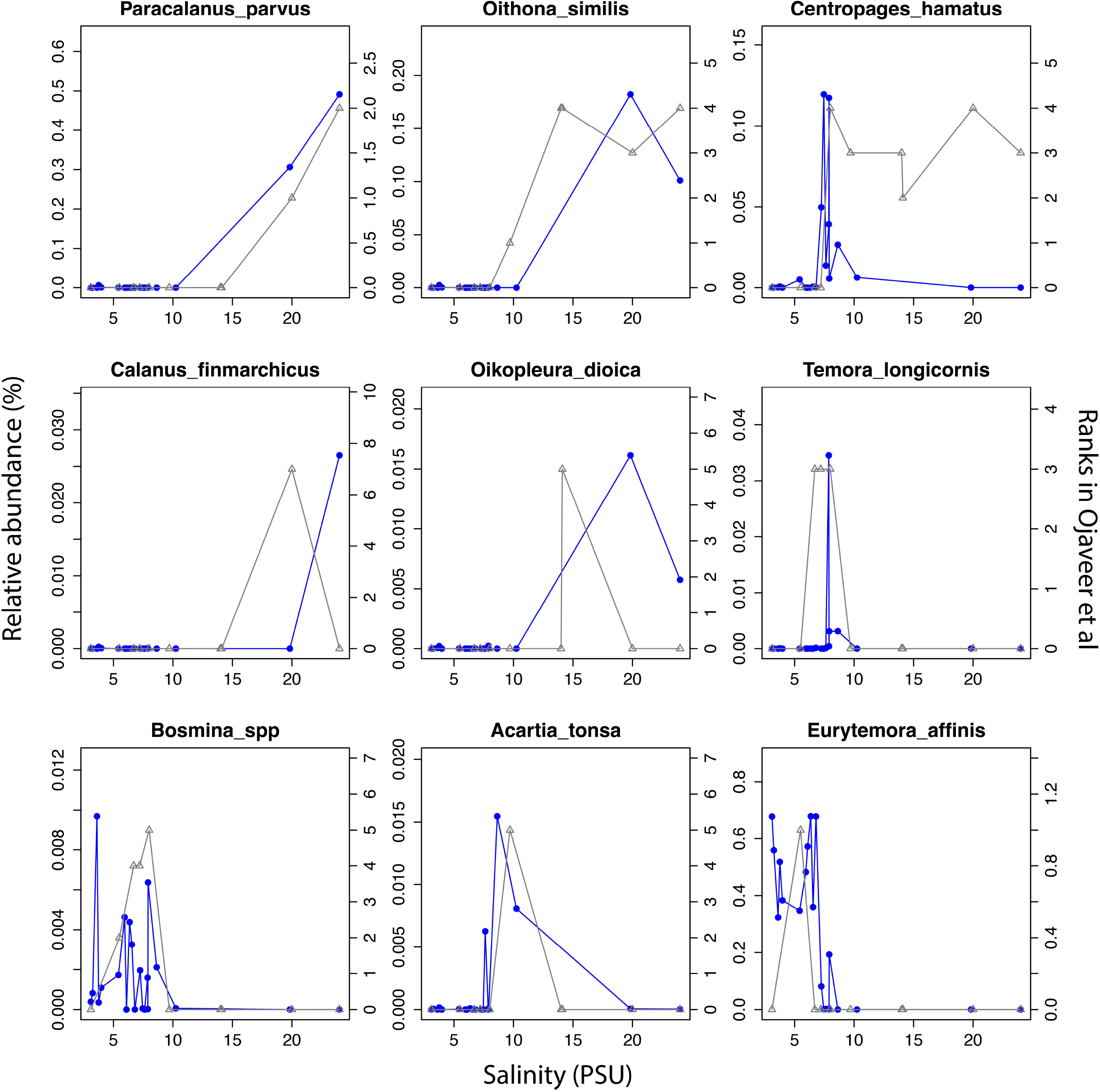
Abundance distribution of dominant mesozooplankton (Metazoa) across the salinity gradient in comparison to historical monitoring data. Data from Figure 1 and Table 3 of Ojaveer et al (2010) is indicated with triangles connected by grey lines (right y-axis). Relative abundances of the OTUs for the corresponding species (when a species corresponds to multiple OTUs these were summed) are indicated with circles connected by blue lines (left y-axis).

### Trends in Alpha-diversity Along the Salinity Gradient

The deep sequencing allowed us to investigate trends in alpha-diversity over the salinity gradient. Since alpha diversity estimates are biased by sequencing depth (Lundin et al., 2012), the same number of reads were subsampled from each sample. No clear trends in species richness (number of observed OTUs) or combined richness-evenness (Shannon-Wiener index) could be observed for neither bacteria or protists at the total community level (Figure 7 A,B). Likewise, when zooming-in on specific eukaryotic phyla (here reads were subsampled from within each phylum), no convincing trends in alpha-diversity were observed.

**Figure 7.**
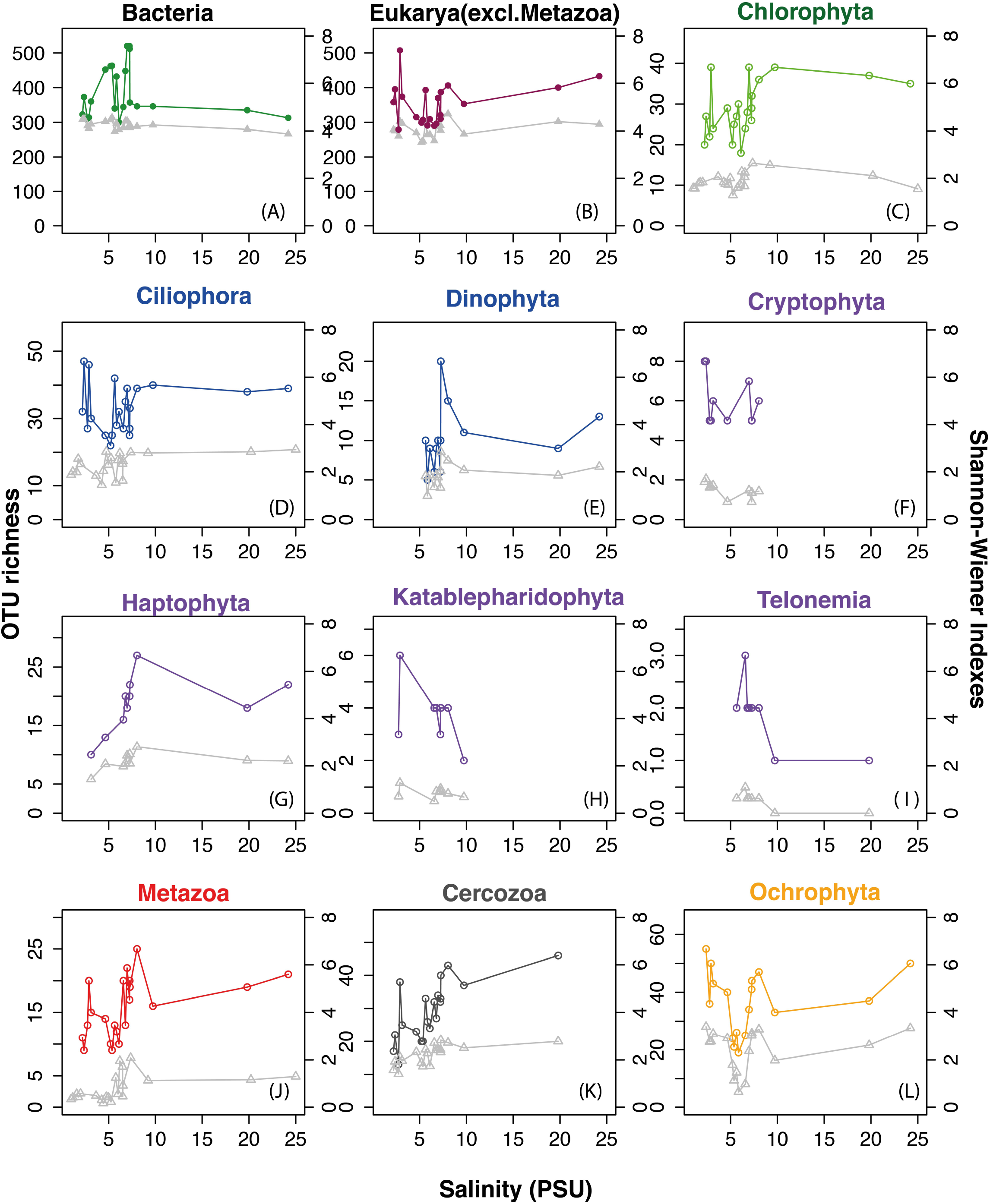
Trends in planktonic alpha-diversity along the salinity gradient. Circles connected by colored lines indicate observed OTU richness (left y-axes) and triangles connected by grey lines represent Shannon-Wiener diversity index (right y-axes). (**A**) Total bacterial community. **(B)** Total protist community (non-metazoan eukaryotes). In order to remove biases due to variation in sequence depth, the same number of reads were subsampled from each sample. In (A) and (B), 11,370 reads were subsampled from each sample. **(C-L)** Eukaryotic phyla with mean relative abundance >1%. For each phylum, the same number of reads were subsampled from each sample. Subsampling was done only from reads belonging to the respective phylum, to avoid that diversity indices were biased by the relative abundance of the phylum within the sample. A minimum of 1000 reads were subsampled per phylum/sample. Samples that contained less than 1000 reads of the phylum were excluded. Phyla were colored according to superphylum in accordance with Figures 4 and 5.

## Discussion

The Baltic Sea has an exceptional long record of plankton monitoring, based primarily on microscopy, from the HELCOM-COMBINE monitoring program (HELCOM, 2015), but to our knowledge this is the first investigation of eukaryotic plankton diversity along the full salinity gradient using high-throughput sequencing. Compared to traditional microscopy based methods, high-throughput sequencing of taxonomic marker genes (metabarcoding) potentially offers a more comprehensive view of microbial diversity. While not yielding absolute cell counts, sequencing data presents integral/structural information of microbial communities, and reveals trends in community changes. While morphological features in some cases can be used to classify plankton with high taxonomic resolution, this is not the case for the majority of species that are too small to allow classification by light microscopy. This is probably the reason why, with only one week’s sampling, the present metabarcoding-based study revealed a microbial diversity on par with what has been recorded in the Baltic Sea with microscopy over a century (Hällfors, 2004; Telesh et al., 2009). In total we observed 1860 OTUs, which can be compared with 2075 known species of phytoplankton (Hällfors, 2004) and 1031 zooplankton (Telesh et al., 2009).

When performing metabarcoding, the taxonomic accuracy is limited primarily by the resolution of the marker gene, the coverage of the primers, and the quality and completeness of the reference database. Here we used the SSU rRNA gene, the most widely used taxonomic marker for bacterioplankton and protists (Amann et al., 1995; Pace, 1997). While it provides a lower taxonomic resolution than e.g. the internal transcribed spacer (ITS) region, its highly conserved regions allow the use of primers with domain-wide taxonomic range. Databases are also more complete than for other markers. We amplified and sequenced 450-600 bp of the hypervariable regions V3-V4 in bacteria and V4-V5 in eukaryotes, using in-house designed primer pairs that cover a large proportion of taxa in each domain (Hugerth et al., 2014a, 2014b). Of the generated 18S OTUs, 31.8% could be classified to the genus level using our automatic classification scheme (see Methods), while 58.3% did not have sufficiently close matches (>98% identity) in the PR2 database (i.e. insufficient database completeness) and 9.8% matched multiple genera (i.e. insufficient taxonomic resolution of the marker). Examples of the latter were *E. Huxleyi* and *G. muellerae* (Bendif et al., 2015) that have 100% identical sequences in the V4-V5 region, as was also the case for species of Labyrinthula isolated from different hosts. Ideally, one would like to sequence both a part of the SSU rRNA gene and a more rapidly evolving marker, such as the ITS. With recently developed emulsion-based protocols, it will be possible to connect different markers from the same cell (Borgström et al., 2015; Spencer et al., 2015).

OTU relative abundances need to be carefully interpreted due to variations in the respective copy numbers of rRNA genes among phylogenetic groups, especially for eukaryotes (Prokopowich et al., 2003; Větrovský and Baldrian, 2013). Due to large cell volumes and vast genomes, dinoflagellates, diatoms and ciliates in general possess more copies of the 18S rRNA gene than smaller flagellates, such as prasinophytes (Godhe et al., 2008; Gong et al., 2013; Zhu et al., 2005). Although previous studies have shown a correlation between cell size (volume or length) and rRNA gene copy number (Godhe et al., 2008), copy numbers can also vary among species of the same genus. Thus, comparing the relative abundance among different taxonomic groups may be misleading. However, the variation in relative abundance of the same group across samples is still valid, allowing to test for correlations with variations in e.g. environmental parameters. Most taxonomic groups, at all taxonomic levels, displayed strikingly smooth abundance changes along the salinity gradient sampled (Figures 2-5, Supplementary figure 5) in contrast to the spiky abundance distributions to be expected if the methodology had introduced much random noise. Systematic biases still likely exist, caused by e.g. primer mismatches and differences in rRNA gene copy numbers, but these should affect all samples equally.

Extracellular DNA released from dead (Choi et al., 2004; Kloos et al., 1994; Rice et al., 2007) or live (Böckelmann et al., 2006) cells may also lead to skewed estimates of abundance and diversity. Furthermore, metabarcoding detects all life forms that carry the marker gene, muddling the distinction between vegetative cells and other life stages. Some species of dinoflagellates, chrysophytes, cryptophytes and haptophytes are known to form resting cysts after blooms (Adam and Mahood, 1981; Jordan and Chamberlain, 1997; Lichtlé, 1980; Lotter et al., 1998; Warns et al., 2012). One example is *Alexandrium*, a toxic genus of dinoflagellates (Teegarden and Cembella, 1996) that sporadically blooms in the Baltic (Kremp et al., 2009; Witek, 2004), for which we observed high abundance (9.1% of protist reads) in one of the Kattegat samples (Supplementary figure 5B). Whether this reflects a bloom is hard to tell, since the sequences may be derived from cysts, though the cysts are also toxic (Oshima et al., 1992).

Similar to what was observed in previous studies (Dupont et al., 2014; Herlemann et al., 2011), bacterial community composition changed gradually along the salinity gradient, and difference in community composition (beta-diversity) was significantly correlated with difference in salinity. Here we showed for the first time a similar pattern for eukaryotic plankton composition using molecular data. The observed pattern can probably also be driven by a combination of other factors such as availability of micronutrients and composition of dissolved organic matter that covary with salinity across the gradient, but it is well established that salinity differences *per se* is a major barrier for species to cross (Logares et al., 2009). A recent study shows that the bacterioplankton inhabiting the Baltic Proper are not locally adapted freshwater or marine populations, but are rather members of a global brackish metacommunity that most likely adapted to brackish conditions before the Baltic Sea was formed (Hugerth et al., 2015). Whether this also holds true for protists inhabiting this sea remains to be investigated. For endosymbiotic protists, isolated from the surrounding water and protected from osmotic stress, the distribution patterns are likely mainly determined by the salinity preference of the host.

The ‘Remane curve’ (Remane, 1934) illustrates how diversity of freshwater, marine and brackish specialist species vary along a salinity gradient. Total diversity is lowest in the horohariculum (5- 8 PSU) because few freshwater and marine species can tolerate this salinity level and relatively few true brackish specialists exist that occupy this niche. Remane based his ‘ Artenminimum’ (species-minimum) concept (Remane and Schlieper, 1971) on macrozoobenthos, but since then the generality of the pattern has been debated (Attrill, 2002; McLusky, 1983; Portillo et al., 2012; Reid, 1961). Recently, Telesh et al suggested the opposite pattern for protists, with a species-maximum in the horohariculum (Telesh et al., 2011b). The analysis was based on pooling species observations within salinity ranges from multiple studies, but has been questioned, since the intermediate salinity where the highest diversity was observed was also the most frequently sampled (Ptacnik et al., 2011; Telesh et al., 2011a), hence the pattern could stem from sampling biases. Here we took the opportunity to measure alpha-diversity with our metabarcoding dataset. This type of data has the potential to give an estimate of diversity that is not biased by morphological characters of the specimens, and the deep sequencing allows subsampling of each sample to the same depth, improving comparisons between samples (Lundin et al., 2012). Similar to what was previously observed (Herlemann et al., 2011), total bacterial alpha-diversity did not display any clear trends over the freshwater-marine continuum. Likewise, the total protist community (eukaryotic data excluding metazoa) displayed neither a species-minimum nor a species-maximum in the horohariculum. There is considerable noise in the data, and the limited number of samples were collected during a single cruise in the summer, thus more data is obviously needed to derive stronger conclusions. However, if pronounced differences in alpha-diversity did exist between salinity regimes, this should have been visible. Telesh attributed the lack of an ‘ Artenminimum’ for protists to their fast evolutionary rate and hence ability to adapt to brackish conditions (as opposed to macroorganisms) (Telesh et al., 2011b), which is also what was suggested for bacteria (Herlemann et al., 2011). However, our recent study that shows that many of the bacterioplankton inhabiting the Baltic Sea are brackish specialists adapted to these conditions before the Baltic Sea was formed (Hugerth et al., 2015), and likely migrated from other brackish waters, opens the possibility that the same applies to some of the protist species. Future population genomic analysis may be able to reveal such patterns.

## Conflict of Interest Statement

The authors declare that the research was conducted in the absence of any commercial or financial relationships that could be construed as a potential conflict of interest.

## Funding

This work was supported by BONUS BLUEPRINT project, supported by BONUS (Art 185), funded jointly by the EU and the Swedish Research Council FORMAS. It is also funded by the Swedish Research Council VR (grant 2011-5689) through a grant to A.F.A. Support from the European Union FP7-project JERICO and the Swedish Research Council VR through the Swedish Lifewatch project is also acknowledged. Y.O.O.H. was supported by a scholarship from the China Scholarship Council (CSC#201206950024).

## Acknowledgments

We are grateful to TransAtlantic AB and the captains and crew of the cargo vessel TransPaper for providing space for the Ferrybox-system and technical assistance during the installation of the system and valuable help during sampling. We are also grateful to Jürg Brendan Logue and Conny Sjöqkvist for insightful comments and suggestions on the manuscript. Sequencing was conducted at the Swedish National Genomics Infrastructure (NGI) at SciLifeLab in Stockholm. Computations were performed on resources provided by the Swedish National Infrastructure (SNIC) through the Uppsala Multidisciplinary Center for Advanced Computational Science (UPPMAX).

## Author Contributions

YH, BK and AA conceived and designed the study. BK performed sampling. YH performed molecular work. YH and AA analysed the data. All authors wrote the paper, and read and approved the final version of the manuscript.

